# LipidSIM: inferring mechanistic lipid biosynthesis perturbations from lipidomics with a flexible, low-parameter, systematic Markov Modeling framework

**DOI:** 10.1101/2023.07.26.550768

**Authors:** Chenguang Liang, Sue Murray, Yang Li, Richard Lee, Audrey Low, Shruti Sasaki, Austin W.T. Chiang, Wen-Jen Lin, Joel Mathews, Will Barnes, Nathan E. Lewis

## Abstract

Lipid metabolism is a complex and dynamic system involving numerous enzymes at the junction of multiple metabolic pathways. Disruption of these pathways leads to systematic dyslipidemia, a hallmark of many pathological developments, such as nonalcoholic steatohepatitis and diabetes. Recent advances in computational tools can provide insights into the dysregulation of lipid biosynthesis, but limitations remain due to the complexity of lipidomic data, limited knowledge of interactions among involved enzymes, and technical challenges in standardizing across different lipid types. In this study, we present a low-parameter, biologically interpretable framework named Lipid Synthesis Investigative Markov model (LipidSIM), which models and predicts the source of perturbations in lipid biosynthesis from lipidomic data. LipidSIM achieves this by accounting for the interdependency between the lipid species via the lipid biosynthesis network and generates testable hypotheses regarding changes in lipid biosynthetic reactions. This feature allows the integration of lipidomics with other omics types, such as transcriptomics, to elucidate the direct driving mechanisms of altered lipidomes due to treatments or disease progression. To demonstrate the value of LipidSIM, we first applied it to hepatic lipidomics following *Keap1* knockdown and found changes in mRNA expression of the lipid pathways were consistent with the LipidSIM-predicted fluxes. Second, we used it to study lipidomic changes following intraperitoneal injection of CCl_4_ to induce fast NAFLD/NASH development and the progression of fibrosis and hepatic cancer. Finally, to show the power of LipidSIM for classifying samples with dyslipidemia, we used a *Dgat2*-knockdown study dataset. Thus, we show that as it demands no *a priori* knowledge of enzyme kinetics, LipidSIM is a valuable and intuitive framework for extracting biological insights from complex lipidomic data.

## Introduction

Lipid metabolism is a highly complex system, characterized by its complicated synthesis processes through the convergence of multiple metabolic pathways. This is accompanied by a variety of enzymes that extend and modify the diverse molecules in the production of lipids (1–4). In Nonalcoholic Steatohepatitis (NASH), for example, disruption of lipid metabolism occurs at different stages, leading to changes in the activity of multiple lipid-related enzymes; however, the underlying mechanisms remain unclear (5–7). Targeting individual enzymes in the lipid metabolism pathway can result in a cascade of effects, impacting not only the biosynthesis of the specific lipids produced by the targeted enzymes, but also a broad range of other lipids that are dependent on shared precursors and interacting enzymes (8–11). Furthermore, each lipid type has unique biophysical properties and can impact diverse cellular functions including signaling, aging, energetics, etc., making the overall response to targeted modulation highly complex (12–14). While targeted lipidomics can provide a snapshot into lipidome changes in a biological sample, they do not provide a comprehensive understanding of the enzymatic and genetic factors driving these changes in the lipid profile (15).

Traditionally, computational models have been employed to analyze the synthesis of diverse metabolites and to decipher the mechanisms underlying the changes in lipid profiles (16–19). However, the accurate modeling of lipid metabolism requires a substantial amount of experimental data to parameterize kinetic models for individual lipid metabolic enzymes, many of which are not well characterized (16,19,20). In contrast, constraint-based, genome-scale models are often limited in their resolution and size, making it difficult to include a large number of specific lipid isomers involved in enzyme and substrate competition (21–24). The development of comprehensive lipid synthesis models is further hindered by the technical challenges and significant effort required for the standardization of multiple lipid species from mass spectrometry (MS)-based lipidomics(25).

Probabilistic modeling (26–28) could now be applied to lipid metabolism without the need for extensive parameterization. This approach is particularly useful given the size, complexity, and incomplete knowledge of lipid metabolism, with its intricate dynamics for the syntheses of fatty acids and glycerol/glycerophospholipids. Here, we developed a Markov modeling approach (Lipid Synthesis Investigative Markov models, or LipidSIM) to study lipid synthesis and degradation, allowing the discovery of perturbations that could lead to observed changes in high-dimension lipidomic data. LipidSIM can flexibly accommodate the stereochemical ambiguity of lipidomic data, and it requires fewer assumptions and parameters when compared to traditional kinetic models(16). The highly modular nature of *de novo* lipid synthesis enables a recursive model construction process to generate a diverse range of lipid species.

We demonstrate the value of this framework LipidSIM with three case studies. First, we analyzed hepatic lipidomic data (*Keap1* dataset) from our own Gubra-Amylin NASH (GAN) diet-fed murine models treated with antisense oligonucleotides (ASO) targeting *Keap1*, a key regulator of cellular oxidative stress implicated in NASH progression (29–31). Through this application, we systematically characterized the perturbed lipidome at different resolutions to generate testable hypotheses regarding lipid biosynthesis reactions. Second, we used a previously published high-dimensional lipidomic dataset from healthy murine models that were repeatedly injected intraperitoneally with CCl_4_ (32,33) to induce fast NAFLD/NASH development and study the progression of fibrosis and hepatic cancer(34–36). This dataset showed how LipidSIM provides biological insights regarding the dosage effect from CCl_4_ and the resulting dyslipidemia. Finally, we applied LipidSIM to our own plasma lipidomic data (*Dgat2* dataset) from lean and GAN diet-fed murine models, treated with a *Dgat2* ASO, which mitigates dyslipidemia in murine NASH models (37–39). In addition to consistency with transcriptomic data, LipidSIM demonstrated the potential for clinical applications by predicting the treatment labels from the model fluxes without requiring cross-sample standardization. In summary, LipidSIM provides a platform for deciphering changes in the lipidome and can integrate the lipidomic and transcriptomic data to decipher indirect and confounding changes in lipid biosynthesis.

## Results

### LipidSIM provides a low-parameter framework for simulating lipid biosynthesis from lipidomic data

We curated a list of lipid biosynthesis reactions in human and mouse (**Supplement 1**) from literature and databases (40–50). The list was used to programmatically generate a general lipid biosynthesis network in the form of a Markov process, resulting in 23,403 stereochemically distinct lipid species (as nodes) and 39,615 synthesis steps (as edges) when untrimmed (see **Methods**). Given a lipidomic dataset, fifty models were fitted per lipid type *k* (e.g., triglyceride [TG], diglyceride [DG], or fatty acids [FA]) per biological sample *j*. Briefly, the lipidomic data were normalized to the total lipid pool of the specific lipid type in the sample (**Figure 1A**). To generate a fitted model, 10^5^ random walks were conducted in the generic network to obtain the frequencies of traversing each edge representing the conversions of different lipid species (**Figure 1B**). The starting point of each random walk was a measured lipid species, with the starting point chosen based on the relative proportion of that lipid species in the sub-lipidome of a lipid type *k* from sample *j* (**Figure 1B-C**). The resulting fitted model (*G*_*j,k,z*_ (*E*_*n*,_ *V*_*m*_)) was composed of all traversed edges and nodes, with weights computed from the frequencies of the random walks (**Figure 1D**). This process was repeated fifty times per lipid type *k* per sample *j*. See methods for details on model construction and fitting.

**Figure 1:**
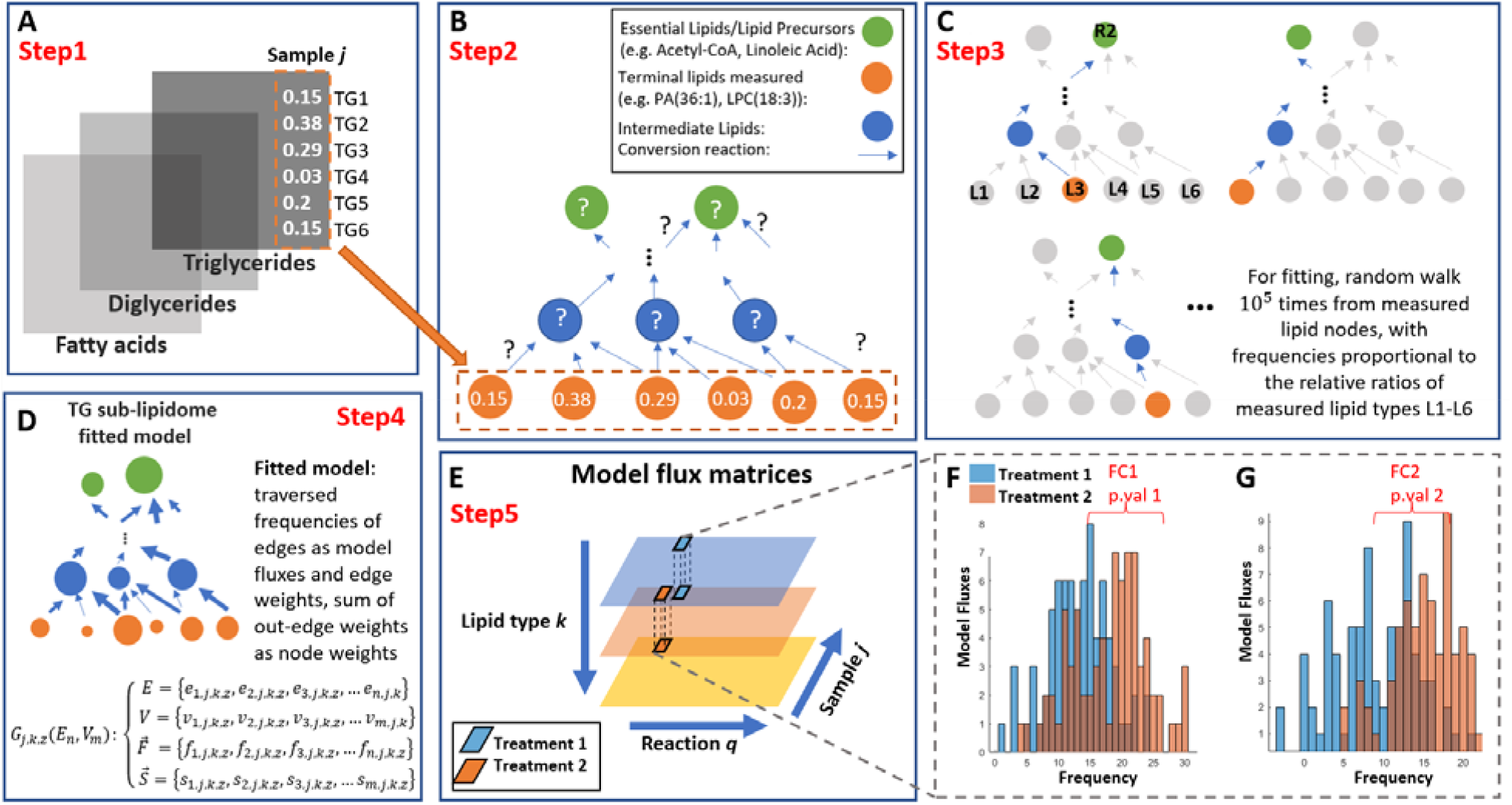
The LipidSIM framework – a schematic overview of the general Markov model fitting process for a lipidomic dataset. **(A)**. For each lipid type k (e.g. triglycerides or TGs, diglycerides or DGs, fatty acids or FAs), the raw MS signals are normalized by the total signals of all the lipid species of lipid type k of each sample j. **(B)**. A fitted model is obtained by random walks from the measured lipid species (leaf nodes) to root lipids (root nodes) given the pre-constructed lipid synthesis network. The starting nodes (lipid species) of the random walks are proportional to the distribution 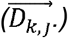 of the measured lipid profile for the lipid type k, sample j. **(C)**. Multiple (10^5^) random walks are conducted to obtain one fitted model from the lipidomic measurement for each lipid type k and of a sample j. Traversed path (without retracing) from a terminal node to a root node is memorized. **(D)**. The total frequencies of traversing edges after all 10^5^ random walks are recorded, resulting in a fitted model (G_j,k,z_ (E_n,_ V_m_)), where E_n_, V_m_ represents the sets of edges and nodes traversed, respectively, with specific edge weights (f_n,j,k,z_) and node weights (s_m,j,k,z_). The fitting process is repeated 50 times per measured lipid type k per sample j. **(E)** Organization of edge weights (pseudo-fluxes) into model flux matrices (MFMs) for the entire lipidomic dataset. To compare model fluxes of a specific reaction between two treatments, model flux vectors could be readily extracted from MFMs (see **Methods). (F-G)** An example comparison of pseudo-fluxes, testing if a reaction differed between two treatment groups. See **Methods** for mathematical details.

After fitting 50 models per lipid type *k* per sample *j*, the model pseudo-fluxes can be organized into model flux matrices (MFMs), allowing systematic differential assessment of reactions between treatments (**Figure 1E-G**). Briefly, when comparing two treatments, a reaction is deemed significantly different if its distributions of model fluxes differ among all models fitted from each lipid type (**Figure 1F-G**). See **Methods** for details on the process of constructing MFMs and calculating relevant statistics. LipidSIM can be used to infer and validate hypotheses regarding changes in the quantities of lipid species (pseudo-concentrations, *s*_*n,j,k,z*_) or reaction fluxes (pseudo-fluxes, *σ*_*m,k,j,z*_) by comparing models fitted from the different lipid types (see **Methods**). By aggregating these hypotheses, a comprehensive understanding of the treatment impact can be gained, shedding light on which enzymatic steps drive the observed differences in the lipidomes.

### LipidSIM elucidated how the KEAP1 ASO modifies the lipidome in GAN-fed murine models

We first demonstrate the basic analysis steps of LipidSIM to study the knockdown of *Keap1* by ASOs using GAN-fed murine models (see **Methods**). The hepatic lipidomes were collected and used for model fitting, including phoshatidylcholine (PC), phosphatidylethanolamine (PE), lysophoshatidylcholine (LPC), lysophosphatidylethanolamine (LPE), and fatty acids (FA) (**Supplemental Data 1**). Direct examination of the lipidomic data comparing treated versus control groups suggested lipids with highly unsaturated acyl chains were the most significantly differentiated lipid class (**Figure 2A**). However, only a few FA species were significantly different after *Keap1* ASO treatment, including reduced DPA but increased EPA and arachidonic acid (**Supplement S2**). Despite the limited number of measured lipid species, the model predicted the average relative quantities (pseudo-concentrations) of all intermediate lipids (i.e., theoretical precursors to the measured lipids) (**Supplement S3**). Analysis of the fitted models revealed significantly higher relative quantities of sn1-LPC(20:4) but lower relative quantities of sn1-LPE(18:2) following *Keap1* knockdown (p-values <1e-3) (**Figure 2C-D**), a result consistent with the actual lipidomic data.

**Figure 2:**
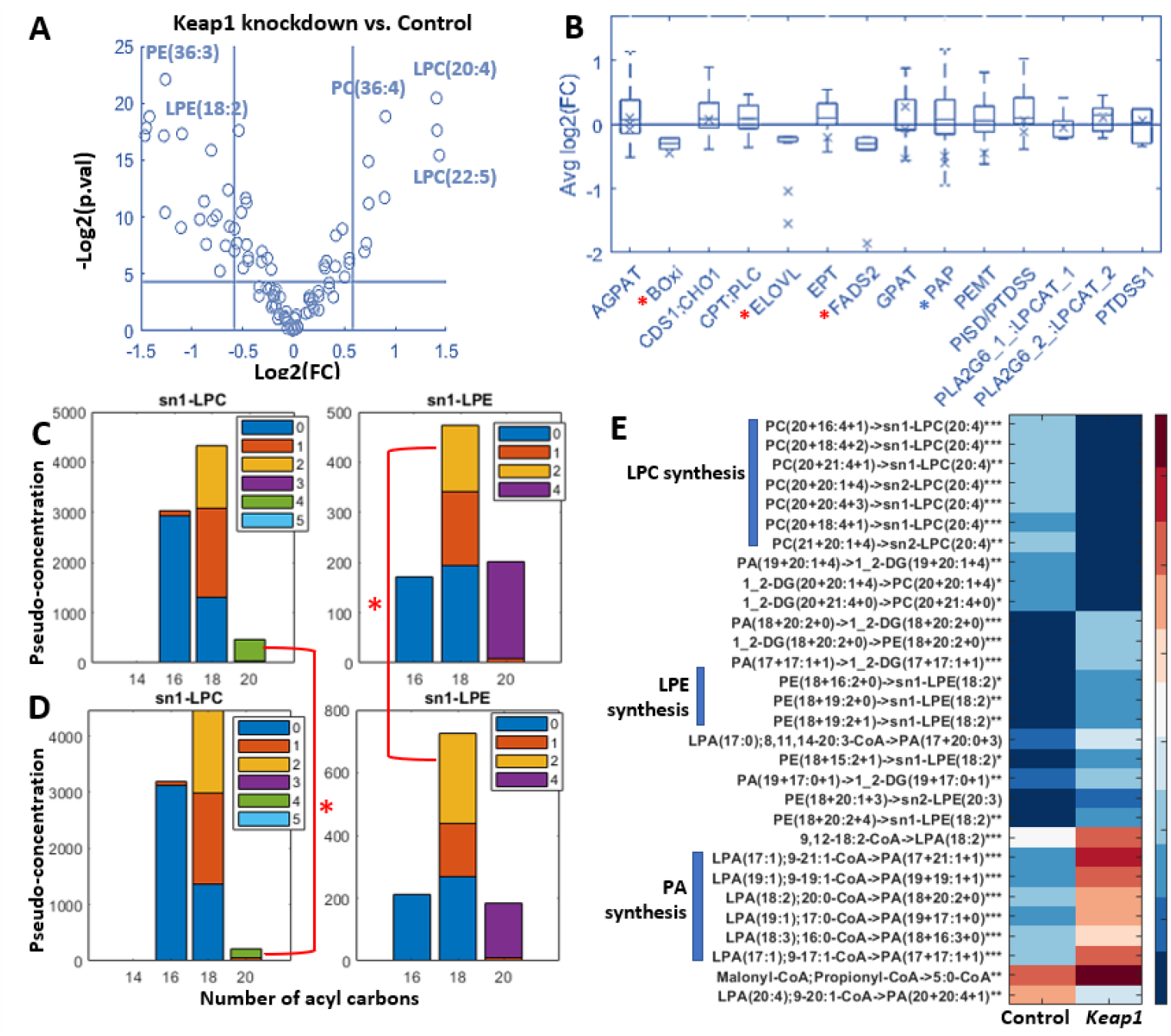
LipidSIM systematically characterized the lipidomes of GAN-fed murine models after KEAP1 knockdown. **(A)**. Volcano plot of differential lipid species from the Keap1 dataset. **(B)**. Boxplot of the distributions of log_2_ fold changes of pseudo-fluxes following Keap1 knockdown, classified by reaction types. The crosses (“x”) indicated the log2 fold changes of related gene levels (CHIP-Seq) previously reported in Keap1^−^/^−^ mice (51). The red asterisks indicated significant differentials of both the model-inferred flux fold changes and the relevant gene levels, whereas the blue asterisks indicated significant differentials for the relevant gene levels or model fluxes only (|log_2_(FC)| > 0.56). **(C-D)**. Aggregated pseudo-concentrations of sn1/2-LPC/LPE lipid species by number of acyl carbons and unsaturated carbon bonds on the acyl chains (represented by different colors). The top two panels were associated with control ASO samples, and the bottom two samples were associated with Keap1 ASO samples. **(E)**. Top significantly perturbed reactions based on the fold changes of the model pseudo-fluxes between the two treatment groups (Keap1 ASO vs. control ASO). (*: p-value <5e-2; **: p-value <1e-2; ***: p-value < 1e-3).

To investigate how changes in model fluxes contributed to the observed lipidome perturbations from the *Keap1* ASO treatment, we calculated the distributions of log_2_ fold changes of pseudo-fluxes (**Figure 2B**). We found that the reaction pseudo-fluxes of beta-oxidation (BOxi), elongation of polyunsaturated very long chain fatty acids (VLCFA [ELOVL]), and desaturation of long chain fatty acids (LCFA [FADS2]) were significantly reduced (see **Methods)**. Encouragingly, the model-predicted changes in reaction flux were consistent with differential expression of relevant genes found previously in *Keap1*^−^/^−^ mice (51), including *Acadl* (BOxi), *Elovl2/5* (ELOVL), and *Fads2* (FADS2) (**Figure 2B**). The significance of model flux fold changes was consistent with the expression changes of genes catalyzing these reactions (**Supplemental Data 2**). Peculiarly, while the previous study also suggested a decrease in *Lipin2* (PAP) expression, the models did not show significantly different changes in PA-to-DG synthesis (PAP) (**Figure 2B**). We anticipate that this increased DG synthesis, a hallmark of steatosis during NASH progression (52,53), might counteract the decrease in *Lipin2* expression seen in normal chow-fed *Keap1*^−^/^−^ murine models. At a finer resolution of model fluxes, the models indicated that LPC(20:4) was the product of the most severely reduced LPC conversion reactions (from PC), and LPE(18:2) was the product of the most increased LPE conversion reactions (from PE) (**Figure 2E**). Interestingly, while the pseudo-fluxes of PA synthesis reactions (GPAT) were not significantly different between the treatment groups as whole (**Figure 2B**), the model fluxes suggested that there was increased PA synthesis reactions involving 18:2/3 and 17/19:1 FAs (**Figure 2E**).

### LipidSIM dissected the mechanisms driving the divergent lipidomes from high-dose CCl_4_

The effects of different doses of CCl_4_ on murine hepatic lipidomes were explored using LipidSIM on a published dataset (33). Briefly, the hepatic lipidomes of fatty acid (FA), phosphatidic acid (PA), diglyceride (DG), and triglyceride (TG) were measured from mice intraperitoneally injected with control (C, olive oil only), 1% (CL), 5% (CM), or 10% (CH) CCl_4_ (10mL/kg of olive oil). While we observed significant depletion of these lipid types in the low and medium CCl_4_ groups (Student’s t-test, p-value <0.01), the high CCl_4_ group showed smaller or insignificant deviations from the control group (**Figure 3A-D**). Despite clear differential clustering of TG and DG species among the control (C), CL/M, and CH samples, a closer examination of the lipid labels of individual blocks failed to provide any clear biological insights for these clusters (**Figure E-F**).

**Figure 3:**
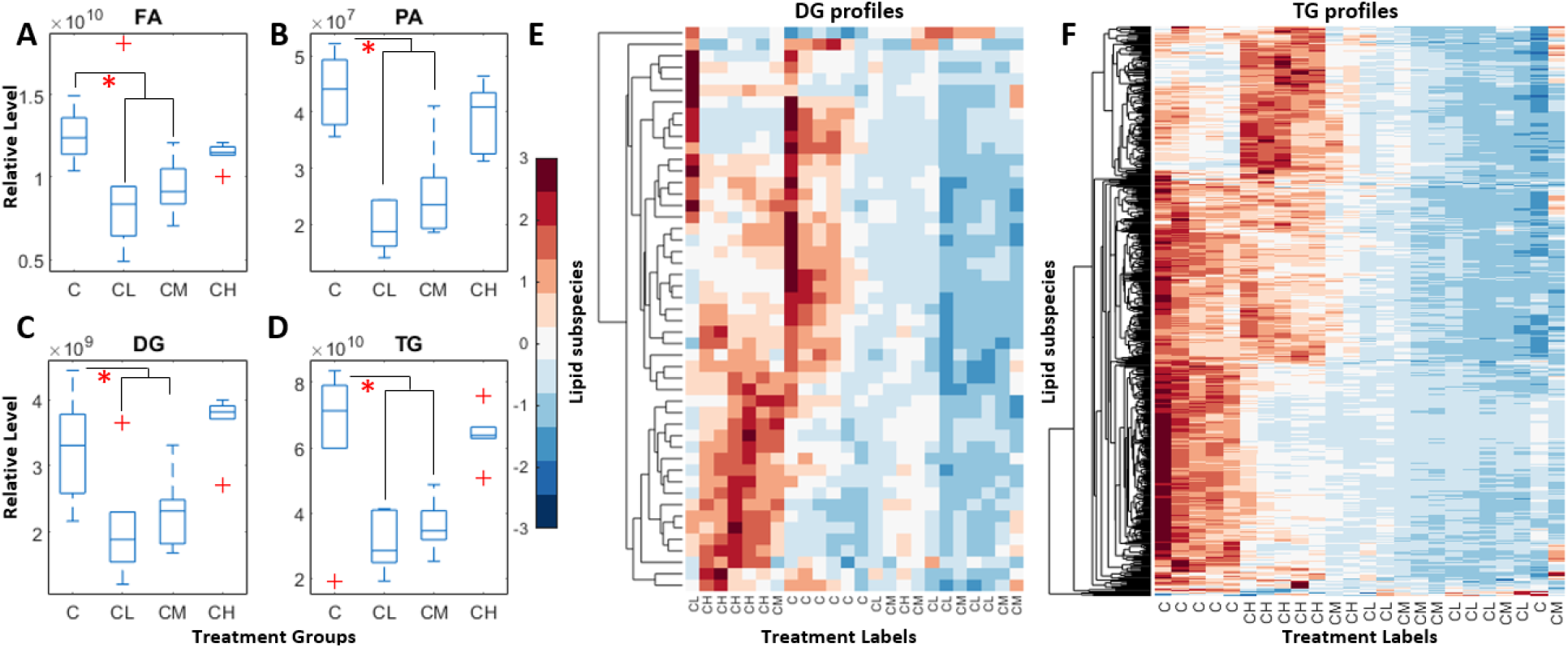
Overview of the raw lipidomic data from mice treated with CCl_4_, starting with a lipid pool overview (A-D) and delving into finer details (E-F). **(A-D**) The total relative lipid pool sizes of different lipid species from four treatment groups (C: control; CH: high dosage CCl_4_; CL: low dosage CCl_4_; CM: medium dosage CCl_4_). Y axis represents the total area of raw MS data and x axis represents the treatment groups. **(E-F**) visualization of the hierarchical clustering of diglyceride (DG)/triglyceride (TG) lipidomic datasets by lipid species and treatment labels, normalized by lipid species (rows).

To understand the differences in the models, we compared fold changes by reaction type between the medium *CCl*_*4*_ dosage (CM) and high *CCl*_*4*_ dosage (CH). The control samples (C) were used as the baseline for comparison. We found that at high CCl4 dosage (CH), the fluxes through lipid synthesis pathways such as LPA/PA/DG/TG synthesis (AGPAT/GPAT/PAP/DGAT2), SFA synthesis, VLCFA desaturase reactions (FADS2), and MUFA desaturase reactions (SCD) were all significantly higher than those at medium CCl4 dosage (CM) (**Figure 4A-B;** see **Methods**). The extent of the increases was substantial that, in the cases of FASN/AGPAT/GPAT/PAP/DGAT2 reactions, reductions of model fluxes in the CM samples (**Figure 4A**, medians below the baseline Avg log_2_(FC) = 0) were reversed in CH samples (**Figure 4B**, medians above the baseline Avg log_2_(FC) = 0). These divergent regulated directions of these reaction fluxes seemed to indicate a shift in lipid homeostasis from the control samples to both CM and CH samples, but in opposite directions.

**Figure 4:**
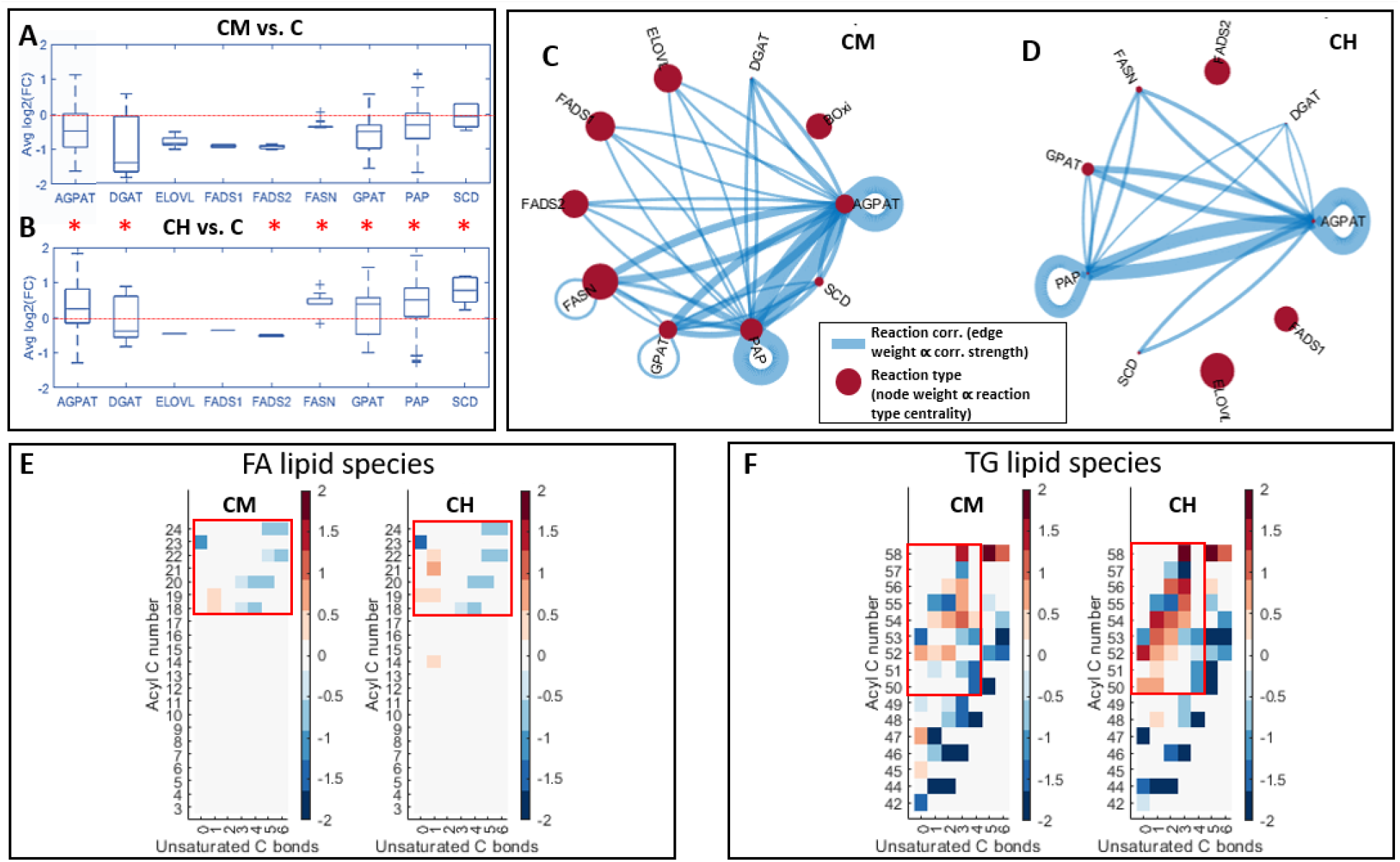
The modeling results revealed perturbations to lipid homeostasis could lead to divergent lipidomes observed in CM and CH samples. **(A-B)**. Boxplot of the distributions of log_2_ fold changes of pseudo-fluxes from control samples to **(A**) CM samples or to **(B**) CH samples by reaction types. The red asterisks indicated significant differentials between the two treatment groups in terms of the fold change distributions (see **Methods** for criteria for significance). (**C-D)**. Correlations between the model flux fold changes for **(C)** CM samples or for **(D)** CH samples by reaction types. The edge width was indicative of a significant correlation between two reaction types, and the node size was indicative of how important the perturbation of the reaction type was predictive of perturbations of other reaction types. (**E-F)**. The heatmaps show fold changes of lipid pseudo-concentrations by lipid species from the control group to a medium (left) or high (right) dosage CCl_4_-treated group. Rows of each heatmap represent total acyl carbon numbers, and columns represent numbers of unsaturated acyl carbon bonds. Only significant fold changes (|log_2_(FC)| > log_2_(1.3), p-values < 0.01) were shown by the heatmaps.

To investigate how lipid homeostasis could be differentially disrupted between CM and CH samples, correlation networks between reaction types were generated using normalized model fluxes (**Figure 4C-D**). The nodes represented distinctive reaction types in the lipid biosynthesis networks and the edges indicated significant correlations between perturbations of these reactions (calculated as fold change in model fluxes from baselines, see **Methods**). The edge width (**Equation 9**) was indicative of a significant correlation between two reaction types, while the node size (**Equation 10**) reflected the importance of the associated reaction type’s perturbations in predicting perturbations of other reaction types. The network generated from CM samples (**Figure 4C**) demonstrated that almost all lipid biosynthesis reaction types were significantly correlated with DG and LPA syntheses, as expected, given that their acyl chain lengths and degree of unsaturation reflected the ratios of FA species in the FA pools, and the products of these reactions were immediate precursors for TG syntheses. However, the network generated from CH samples (**Figure 4D**) showed a loss of significant correlations between the VLCFA/PUFA synthesis reactions (FADS2, FADS1, ELOVL) and all the subsequent reactions incorporating their acyl-chain products (AGPAT, GPAT, PAP). The large relative centralities of FADS2, FADS1, and ELOVL, combined with the fact that they lacked correlation with other reaction types, implied their fluxes as bottlenecks (i.e., high centralities) for very few subsequent reactions (i.e., insignificant correlations).

Corroboratively, we performed a visual analysis of the lipid type with the most perturbed model pseudo-concentrations (see **Methods**). The results revealed that the high CCl_4_ dosage (CH) had a significant increase of TG species with limited unsaturation of acyl carbon bonds (0-3) and large acyl carbon numbers (50-58) (**Figure 4F**), but such dramatic increase was not observed in the FA species between the CM and CH groups (**Figure 4E**). This observation, in conjunction with the correlation analyses, suggests that the synthesis of VLCFA/PUFA as reaction flux bottlenecks may be resulted from the elevated synthesis of glycerolphospholipids at a higher CCl_4_ dosage (in CH samples).

### LipidSIM reconciled lipidomic and transcriptomic changes following Dgat2 ASO treatment

Using the *Dgat2* dataset, we demonstrated the ability of LipidSIM to evaluate the theoretical effect of *Dgat2* ASO on the dyslipidemia of murine plasma lipidome from a longitudinal study with a multivariate experimental design. Briefly, 6-week-old C57BL/6 mice were fed Gubra-Amylin NASH (GAN) diet for 16 weeks before administration of 10mg/kg/week of *Dgat2* (n=8) or control ASOs (n=8) (see **Methods**). Plasma lipidomes were collected from age-matched lean mice (n=4), mice fed with GAN diet that were administered control ASO (n=8) or *Dgat2* ASO (n=8), at weeks 1 and 4 of the drug administration schedule (see **Methods**). Additionally, the transcriptomics (Digital Gene Expression-DGE) data were collected and processed from the *Dgat2* samples at the same time points (see **Methods**).

The analysis of the aggregated lipidomes revealed the challenges of accurately determining the effect of treatment in the absence of longitudinal data and in the presence of mice on different diets (e.g., lean and GAN-fed groups). Notably, the comparison of the lipidomes between the control and *Dgat2* samples was made difficult by the fact that they started with different lipid pool sizes at week one (**Figure 5A-C**). Furthermore, the existence of outliers (**Figure 5B-C**) despite the small sample sizes and the large variances of the lean FA/DG pools at week 4 (**Figure 5A-B**), raised concerns regarding batch effects and limited the power of statistical tests with the raw lipidomes.

**Figure 5:**
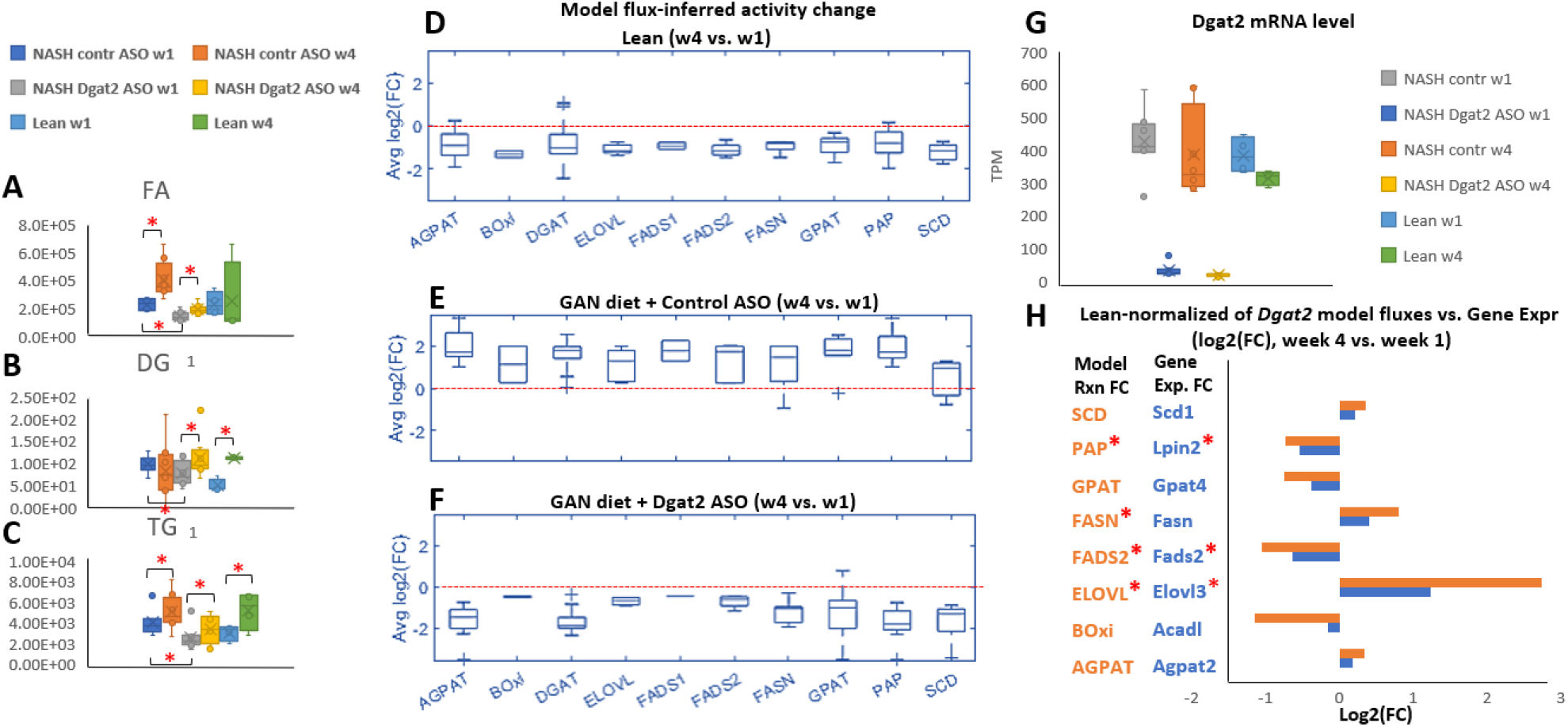
Aggregated relative quantity of lipid pools of (A). FA, (B). DG, and (C). TG from the Dgat2 lipidomic dataset. **(D-F)**. Boxplot of the distributions of log_2_ fold changes of pseudo-fluxes from week 1 to week 4 from **(D)** lean samples, **(E)** control samples, and **(F)** Dgat2 samples. **(G)**. Dgat2 mRNA levels of all the treatment groups, at week 1 and week 4. **(H)**. log2(fold changes) of model fluxes and associated key gene (average) expression from week 1 to week 4. By regarding lean samples as the baseline lipid homeostasis, the model fluxes were first normalized by the lean sample fluxes at the corresponding time points before the computation of temporal fold changes. The red asterisks indicated the significance of fold changes for either gene expressions or model fluxes.

Despite the challenge of deconvolving the lipidomic data, inspecting temporal fold changes of the fitted model reaction fluxes revealed the significant impact of *Dgat2* ASO treatment on many lipid synthesis reactions beyond *Dgat2* knockdown (**Figure 5G**). Similar to the lean samples (**Figure 5D**), the *Dgat2* ASO-treated samples displayed significantly reduced fluxes of AGPAT, BOxi, DGAT, ELOVL, FADS2, FASN, GPAT, PAP, SCD, with the exception of FADS1 (**Figure 5F**), whereas the control ASO-treated samples showed significantly increased fluxes of all the aforementioned reactions (**Figure 5E**). The resemblance of systematic lipid synthesis reduction over time between the *Dgat2* and the lean samples but not between the control and the lean samples supported the strong corrective effect of *Dgat2* ASO on dyslipidemia in the GAN-fed murine models. To validate the impact of *Dgat2* ASO, as inferred from the modeling results, we further examined the expression levels of key enzymes associated with the model fluxes. Using the lean samples as the baseline lipid homeostasis, the model fluxes from the *Dgat2* samples were normalized by the model fluxes of lean samples, at week 1 and week 4, respectively (the procedure is identical as computing the flux fold changes between the *Dgat2* and lean samples, see **Methods**). The temporal (week 4 vs. week 1) fold changes of normalized reaction fluxes for each reaction type were then compared with the temporal fold changes in the expression levels of their corresponding key isozymes (**Figure 5H**). Interestingly, the comparison showed high consistency in the direction of changes between the transcriptomic data and model fluxes, while the magnitudes of the fold changes differed (**Figure 5H, Supplemental Data 3**). The model also accurately captured the significance of perturbations to FADS2, ELOVL, and PAP reactions, whose associated major enzymes also showed significantly differentiated expression levels, except for FASN (**Figure 5H**). Despite the insignificance for FASN flux increase, however, the trend agreed with the expression of *Fasn* and could be an indication of perturbed exogenous sources of saturated FAs feeding into the liver.

### LipidSIM enhances classification of samples using intuitive ensemble decision trees

We tested if LipidSIM could classify and predict the treatment status of samples based on plasma lipidome measurements. For this, we computed the aggregated model pseudo-fluxes for each reaction and sample in the *Dgat2* plasma dataset (see **Methods**). The process to train a decision tree was as follows: the biological samples were randomly partitioned into a training set (75% of samples) and a testing set (25% of samples), with the corresponding model fluxes being partitioned accordingly (**Figure 6A, Step 1**). The Minimum Redundancy Maximum Relevance (MRMR) Algorithm was deployed to filter the features from the training set based on the MRMR scoring metric (**Figure 6A, Step 2**). A binary decision tree was trained using the training set until a misclassification rate of less than 10% was achieved (**Figure 6A, Step 3**). The predicted label for each biological sample was assigned based on the majority label predicted by the 50 corresponding models (**Figure 6A, Step 4**).

**Figure 6:**
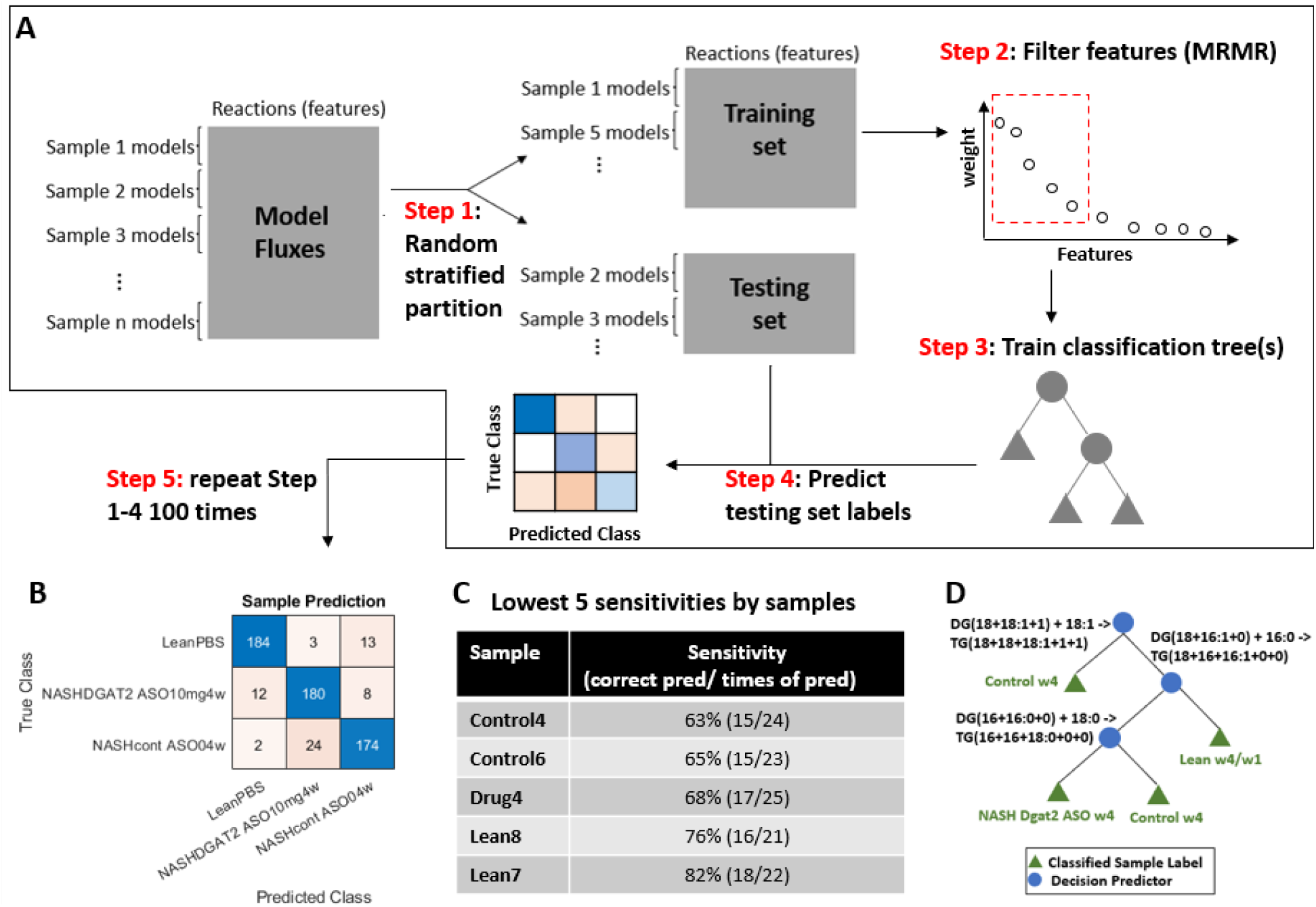
General procedure of fitting a binary decision tree to a training set of aggregated model fluxes and predicting the treatment labels of biological samples. **(A)**. (Step1) Biological samples were randomly partitioned into a training set and a testing set. The multiple sets of model fluxes associated with the biological samples were partitioned based on the biological sample partition. (Step 2) Model features were filtered based on the scoring of Minimum Redundancy Maximum Relevance algorithm prior to model fitting. (Step 3) A binary decision tree was fitted to the training set. A new decision tree would be trained if the misclassification rate was >10%. (Step 4) The trained classification tree was used to predict the treatment labels of the models. The predicted label of a biological sample was determined by the most frequently predicted label of its associated models. (Step 5) **(B)**. Confusion table of true labels vs predicted labels for 100 times of prediction results from randomly sampled training sets and testing sets. **(C)**. Lowest 5 sensitivities by samples. **(D)**. A simple classification tree constructed from a list of top split predictors (**Supplement S4**). The classification tree was able to correctly predict the labels of all the samples and at least 98% of their associated models.

The performance of the machine learning method was tested by repeating steps 1-4 of the method 100 times with randomly selected training and testing sets. The treatment labels of the biological samples were accurately predicted with sensitivity of 0.92, 0.90, and 0.87 for lean, drug-treated, and control sample, respectively; and the specificity of 0.97, 0.93, and 0.95 for lean, drug-treated, and control samples, respectively (**Figure 6B**). A closer examination of the five worst misclassified samples showed that the ensembles of 100 classification trees still usually predicted the treatment labels of these samples correctly (>63% correct prediction) when trained with different training sets (**Figure 6C**). Finally, the most frequently selected split predictors (**Supplement S4**) from the ensemble of 100 classification trees suggested that three predictors involving TG synthesis were able to correctly predict the treatment labels for all samples (**Figure 6D**).

## Discussion

To obtain biological insights from lipidomic data, standard analyses can examine lipid signatures indicative of changes in lipid metabolism(1,54). Alternatively, machine learning methods can associate lipidomic features and other omics or phenotypes, but there is an intrinsic limit in such methodologies to explain which enzymes are involved (15,55,56). To increase the explainability of such analyses, LipidSIM accounts for the lipid biosynthesis network; furthermore, this accounts for the interdependency of all measured lipid species in the lipidomic datasets. For instance, lipidomics of triglycerides and their acyl chains may also contain information regarding the relative quantities and types of fatty acids, which are often measured separately. By introducing the network, LipidSIM effectively explores the possible reaction “fluxes” that achieve the observed lipidome, including all measured lipids. Indeed, it can account for the synthesis of shared precursors across many lipid types. For example, PA is a shared precursor for PE and PC syntheses, and synthesis of PA can therefore be inferred from both PE and PC measurements using the models. When comparing two samples, the range of possible fluxes can be compared statistically, and hypotheses can therefore be generated based on significantly different fluxes between the samples (see **Methods, Figure 1E-G**). Intuitively, the differential reaction fluxes between samples could be readily compared with the corresponding gene expression or protein abundance levels, and the consistency of such comparisons have been repeatedly demonstrated in this study (**Figure 2B, Figure 4A-B, Figure 5D-F**).

### LipidSIM provides comparison of lipidomes at different resolutions without the need of cross-sample normalization

Analyzing large lipidomic data sets remains daunting due to challenges from technical variation and issues with standardizing across samples or lipid types (15,57,58), as observed from the *Dgat2* ASO dataset (**Figure 5A-C**). The sources of the error may include variability in tissue sampling for biopsies and the short half-lives of many derivatized lipid species during mass spectrometry measurements. Thus, the power of statistical tests when directly applied to lipidomes is likely diminished, especially when comparing specific lipid species between treatment groups of multiple samples. LipidSIM addresses this issue by normalizing the total lipid pools by lipid types and by individual samples. Instead of probing the differences of absolute lipid quantifications across samples, LipidSIM investigates the changes of relative ratios of individual lipid species in a normalized sub-lipidome and the relative shift in model fluxes to achieve such observed ratios (e.g., **Figure 4E-F**). It also probes the shifted lipid homeostasis by investigating the changed correlations between theoretical reaction fluxes from one sample to another (e.g., **Figure 4C-D**). At a finer resolution, LipidSIM can also project the theoretical impact of each individual reaction to the overall changes in the observed lipidomes (**Figure 2C-E**). By investigating three distinct lipidomic datasets from both longitudinal and single-timepoint studies, we demonstrated the flexibility and consistency of conducting these differential analyses at different scales systematically using LipidSIM.

Additionally, the simple and highly interpretable binary decision trees showed excellent performance in predicting the treatment labels from the *Dgat2* plasma dataset even when the sample size was very small (6 for each class in a training set) (**Figure 6**), further attesting to the consistency of the model framework. In combination with the fact that cross-sample normalization was not required for model construction, LipidSIM revealed its potential in clinical applications when deployed with machine learning, such as distinguishing treatment phenotypes of diseased patients without the need to collect reference samples, especially for longitudinal monitoring.

### LipidSIM is modular and expandable to account for new knowledge regarding lipid metabolism

Unlike kinetic models or constraint-based models (16,19,59,60), the only input required for LipidSIM is lipidomic data. Also, parameter tuning is not required. The generation of the generic network depends on the user-defined reaction rules, and the models can be modified or expanded when new knowledge regarding these reactions becomes available. Here, we curated a list of common lipid synthesis/degradation reactions commonly seen in humans and mice (**Supplement S1**). The modeling framework automatically trims out unused reactions to improve the power of statistical tests by reducing both the noise and the dimensions of the fitted models. The trimming also benefits machine learning with model parameters as it reduces the risk of overfitting. The model modularity also enables easy grouping of reaction fluxes into different reaction types by desired features, such as the number of acyl carbons and the position of desaturation (e.g., FADS1 vs. FADS2), incorporating knowledge regarding isozyme specificities as it becomes available.

### LipidSIM is well suited to study lipidomes altered by NASH progression and potential treatment impact

To date, while severe deregulation of phospholipid syntheses and glycerolipid metabolism were observed in NASH patients and murine models, the mechanisms driving this remain scattered (61–64). Due to dramatically increased *de novo* lipogenesis, earlier stages of NASH induced by high-fat diet can still ramp up esterification of fatty acids into TG to ameliorate cytotoxicity from free fatty acids and consume DG, but that ability could be compromised with severe fibrosis or necrosis (65). Here we analyzed data on its progression using LipidSIM to quantify the shift of lipid homeostasis, yielding novel hypotheses explaining the divergent lipidome observed from mice injected with high-dosage CCl_4_.

Furthermore, in the other two examples, we found that the models quantified the impact of *Keap1/Dgat2* ASO treatment on the observed dyslipidemia in the murine models. Knocking down either *Keap1*or *Dgat2* mitigates NAFLD/NASH lipid accumulation (29,31,37,38), but the mechanisms and the potential interactions between the lipid biosynthesis enzymes remained largely unclear. In both cases, we showed how lipidomics can be corroborated by transcriptomic evidence by cross-validating the changes of the model-inferred reaction fluxes and the changes of the associated gene expression levels (**Figure 2B, 5H**). In the lipidomic analysis following *Dgat2* ASO treatment, LipidSIM provided insights into the treatment’s corrective effect (**Figure 5F**) by comparing the lean and the drug-treated samples, despite of the large within-group variation observed from raw lipidomes (**Figure 5A-C**). This intuitive, versatile modeling framework is thus useful in studying the deregulated lipidomes in metabolic diseases such as NASH, and the potential genetic interactions among the many lipid synthesis isozymes via knockdown experiments.

## Acknowledgements

This work was supported by generous funding from Ionis Pharmaceuticals and NIGMS (R35 GM119850). Lipid analysis was performed at the UCSD Lipidomics Core.

## Methods

### Data Preprocessing

To ensure completeness, consistency, and metadata accuracy, the *Dgat2* plasma lipidomic datasets of 72 samples (lean wildtype, NASH + control ASO, NASH + *Dgat2* ASO 10 mg/kg, at week 1 & week 4) and *Keap1* lipidomic datasets of 16 samples (NASH + control ASO, NASH + *Keap1* ASO, both at week 2) were curated, and the total relative intensities by lipid types were visualized by bar plots. The progression dataset (raw MS signal area) and metadata was retrieved from Metabolomics Workbench (Study ID: ST002097) and included the sample IDs SA201216-SA201239. For each lipid species measured, the relative intensities across samples and lipid subspecies were normalized by a global standard used during the MS analysis. For model fitting, we normalized the total lipid pool size to be 1 for each sub-lipidome of lipid type *k* per sample *j*. The normalized distribution was represented by a vector 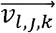, where *l* denoted a specific lipid specie (e.g. TG(18+18+18:0+0+1)) of lipid type *k* from sample *j*.

### Model Construction & Fitting

The generic lipid synthesis network was generated by a recursive process of converting essential lipids (linoleic acid and alpha-linoleic acid), cholesterol, malonyl-CoA, propionyl-CoA, and acetyl-CoA to other commonly observed or measured lipid species. Currently, the modeling network accounts for syntheses of common fatty acids (saturated & mono/poly-unsaturated), phospholipids (DG, TG, PA, PC, PI, PS, PE, PG, plasmalogens, and plamanyl-PL), and eicosanoids (PG1/2/3). The reaction rules used for the recursive expansion of lipid network were tabulated in **Supplement 1**. The reaction rules were curated from a compilation of literature sources, Uniport, and KEGG Pathway(11,40,41,44–46,66,67). The scopes of the studied lipidomic datasets excluded consideration of PI, PS, PG, plasmalogens, plamanyl-PL and eicosanoids.

To fit a model, the edges of the generic lipid biosynthetic network were reversed and allowed the fluxes of measured lipids (nodes) to travel through the network via all precursing intermediate lipid species until reaching the roots of the network (acetyl-CoA, acetoacetyl-CoA, malonyl-CoA, propionyl-CoA, 9,12-18:2-CoA,9,12,15-18:3-CoA, cholesterol). The generic reversed model *G*(*E, V*) can be described as a set of n edges (*E*) and m nodes (*V*) with edge weights 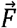 and node weights 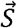. A fitted model (numbered as *z*) for a sub-lipidome of lipid type *k* and sample *j* could therefore be described by **Equation 1**:

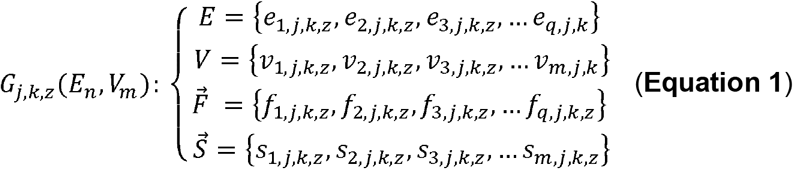

Specifically, 10^5^ random walks were executed to obtain the edge weights and node weights. For each random walk, the starting nodes were chosen sequentially from the set *L*_*l,j,k*_ (**Equation 2**), where the number of times metabolite *l* was chosen as the starting node was proportional to its measured ratio in the sub-lipidome of lipid type *k* from sample *j*:

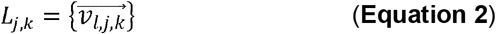

The process of fitting one model can be described by the following pseudocode:

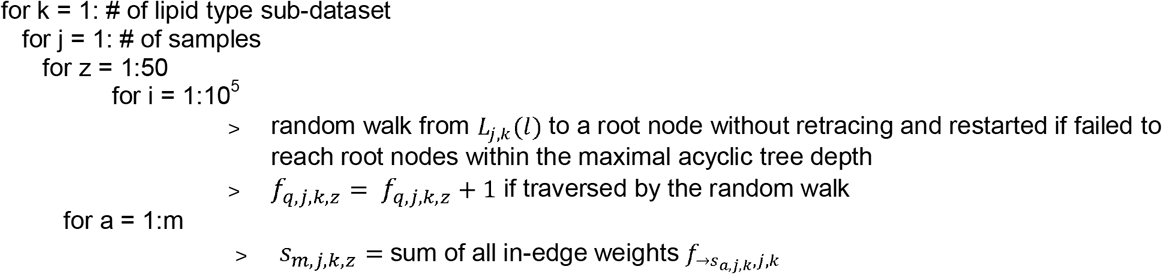

A set of 50 fitted models was generated by the code per lipid type *k* per sample *j*.

For each fitted model, a model pseudo-flux (*σ*_*n,j,k,z*_) was computed as the summation of all edge weights (*f*_*q,j,k,z*_) associated with the reaction *n*, and a model pseudo-concentration (*s*_*m,j,k,z*_) was computed as the summation of the in-edges’ weights. For ease of systematically conducting statistical tests and other manipulations, all fitted model pseudo-fluxes and pseudo-concentrations from a lipidomic dataset were reorganized into model pseudo-concentration matrices (***MCM***_***k***,***j***_) and model pseudo-flux matrices (***MFM***_***k***,***j***_), respectively (**Equation 3**). One *MCM*_*k,j*_ and one were generated per sub-lipidome of lipid type *k* per sample *j*.

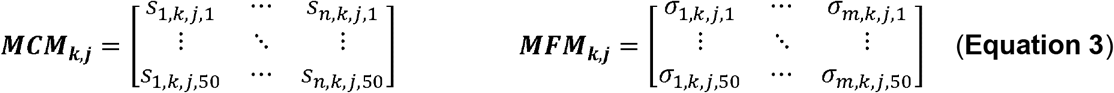

### Computing fold change and statistics of model pseudo-fluxes between two treatments

To compare the models fluxes of a specific reaction *m*_1_ between two treatments A and B, given a specific sub-lipidome *k*_1_, the fold change of models fluxes from A to B was defined as:

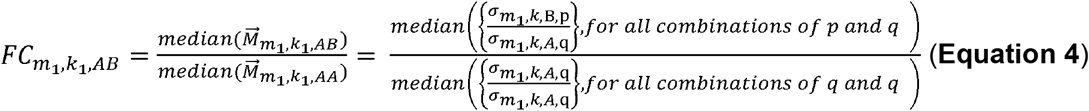

where *p* represented the row indices associated with sample B from ***MFM***_***k***,***B***_ and *q* represented the row indices associated with sample A from ***MFM***_***k***,***A***_. The p-values of the 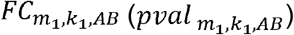 was computed by one-tailed Wilcoxon rank sum test (unpaired) between 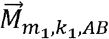 and 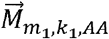 and corrected by Bonferroni procedure when multiple reactions were compared simultaneously.

A reaction was deemed significantly differentiated if the following criteria was met:

a. 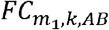 were all >1 or <1 between the compared groups of fitted models, regardless of the sub-lipidome used for fitting (all values of *k*) and except for those carrying no flux through reaction *m*_1._
b. 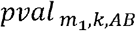 were all smaller than 1e-2 for all compared groups of fitted models, regardless of the sub-lipidome used for fitting (all values of *k*) and except for those carrying no flux through reaction *m*_*1*._
c. The mean of 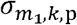 and the mean of 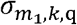 were bigger than 1.

The aggregated fold change of reaction *m*_1_ between treatment A and B was computed as:

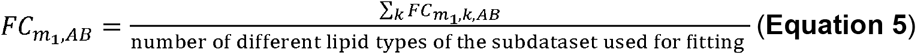

When comparing the differentials of a type of reaction *d* (e.g. all of those catalyzed by the enzyme *Fasn*), the fold change of the reaction type *d* (*FC*_*d,AB*_) was the median of the set 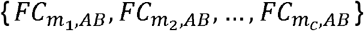, where {*m*_1,_ *m*_2,_ …, *m*_*c*_} were all reactions associated with the reaction type *d*. The significance of *FC*_*d,AB*_ (*pval*_*d,AB*_) was computed by rank sum test between the set of 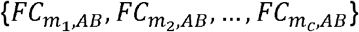 and the set of 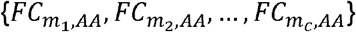, and the threshold was set to be 1e-2.

### Computing fold change and statistics of model pseudo-concentrations between two treatments

To calculate the model pseudo-concentration 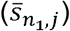 of a lipid specie (or a group of lipid species) *n*_1_ for a sample *j*:

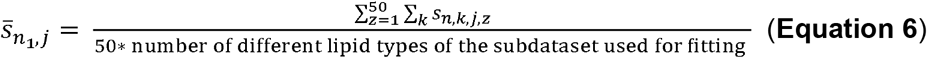

To calculate the aggregate fold change 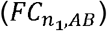 of the pseudo-concentration 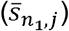 of a lipid specie (or a group of lipid species) between treatment A and B:

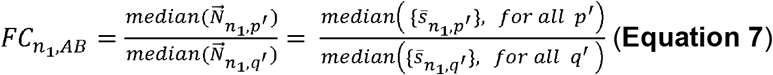

where *p*^’^ represented the row indices associated with sample B from ***MCM***_***k***,***B***_ and *q*^’^ represented the row indices associated with sample A from ***MCM***_***k***,***A***_. The p-values of 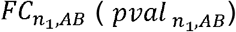 was computed by one-tailed Wilcoxon rank sum test (unpaired) between 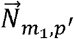 and 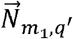 and corrected by Bonferroni procedure when multiple metabolites were compared simultaneously.

### Computing correlations between reaction types and the centralities of reaction types

To compute the correlations between a reaction *m*_1_ and a reaction *m*_2_, fold changes of model fluxes in a treatment group A from treatment group *C* were used as a normalization scheme. Kendall’s 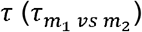 and associated p-values 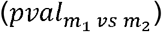 were computed such that:

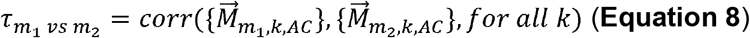

To compute the correlation between a reaction type *d1* and a reaction type *d2*, the correlation metric was defined as:

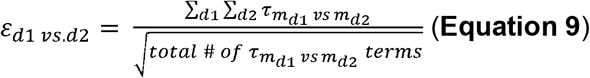

where *m*_*d*1_ and *m*_*d*2_ represented reactions associated with reaction type *d1* and reaction type *d2*, respectively. Only *τ*>0.5 with associated <0.05 were included in the calculation by **Equation 9**.

To understand whether the perturbations to model fluxes of a reaction type *d1* was highly indicative of the perturbations to the model fluxes of other reaction types, an adjacency matrix of all reactions (*m*) was constructed with the significant Kendall’s *τ* as elements (only include *τ* >0.5 with associated *pval*<0.05). Eigenvector centrality (ℂ*m*) weighted by Kendall’s *τ* was computed for each reaction from the adjacency matrix by the *centrality* function in MATLAB. Finally, we define the aggregated centrality (∅) of reaction type *d1* as the mean of all such ℂ*m*:

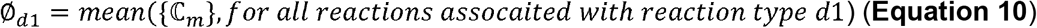

### Experimental designs for KEAP1/Dgat2 ASO-treated mice fed on GAN diet

#### Keap1 ASO experimental design

6-week-old C57BL/6 male mice were fed Gubra-Amylin NASH (GAN) diet (40% high-fat, 20% high-fructose, and 2% high-cholesterol, Cat# D09100301, Research Diets, NJ USA) research diets for 25 weeks before administration of *KEAP1* (n=8) or control ASOs (n=8). The mice were randomized based on body weight and ALT values (Clinical Analyzer AU480, Beckman Coulter, CA USA) before ASOs were administered. Briefly, the mice were administered one dose of Keap1 ASO on week 25, week 26 and week 27 at 2.5mg/kg/week (3 dosages in total) while remaining on the GAN diet. Terminal liver tissue was collected 72 hours following the last dose.

#### Dgat2 ASO experimental design

6-week-old C57BL/6 male mice were fed Gubra-Amylin NASH (GAN) diet for 16 weeks before administration of *Dgat2* (n=4) or control ASOs (n=4). The mice were dosed on week 17 @ 10mg/kg/week and necropsied 48 hours later after the last dosage at 1 week or 4 weeks later (mice were dosed at week 18, 19, 20 and 21 at 10mg/kg/week if not necropsied). Terminal plasma samples and liver tissue were collected at the end of the studies. Liver samples were purified for RNA and submitted for Digital Gene Expression.

### Plasma and hepatic sample preparation for lipidomics and/or RNASeq

#### Keap1 ASO samples

For lipidomics, 20-50mg aliquots of frozen liver tissue were extracted with identical solution with internal standards. 1.7ml of cold extraction solution was added to each tissue aliquot in a 2ml screw top polypropylene tube pre-loaded with 6 ceramic beads (Omni international, Inc. #19-646) and homogenized using the Omni Bead ruptor bead mill (3 × 20 seconds at speed level 5) at 4°C. Samples were deproteinized by incubation in −20°C for 1hr, followed by centrifugation for 15 minutes at 16000g. Supernatant was removed and diluted 1/50 as above for phospholipid runs. 1ml of supernatant was concentrated 3-fold using speed vac and loaded onto PUFA run in a final concentration of 80% Methanol/100uM BHT.

#### Dgat2 ASO samples

For lipidomics, plasma samples were extracted using a modified BUME method(68) and an Equisplash mix (Avanti) containing deuterated lipids was added to 20uL plasma. The extracts were brought to dryness and reconstituted in Buffer (18/1/1 isopropanol/dichloromethane/methanol) and used for LC-MS analysis of PUFA/MUFA and glycerolphospholipids. This process was modified from a previous study(69). For RNASeq, RNA purification was performed on a 6 mm liver biopsy was homogenized in 2 mL guanidine isothiocyanate (Life Technologies, Carlsbad, CA, USA) containing 8% β-mercaptoethanol (Sigma Aldrich, St. Louis, MO, USA). The resulting lysate (30 μL) was applied to the PureLink Pro 96 Total RNA Purification Kit (LifeTechnologies, Carlsbad, CA).

### LC-MS analysis of PUFA/MUFA in Keap1 ASO samples

A semi-quantitative MRM method was used for polyunsaturated (PUFA) and monounsaturated fatty acids (MUFA) measurement, utilizing an API 4500 triple quadrupole mass spectrometer (SCIEX Corp) coupled to a Shimadzu Prominence UFLC (Shimadzu Corp). The method was run in polarity switching mode, collecting half of the 1 sec total cycle time in positive ionizaton mode, then switching to negative mode for the other half. A C18 X-bridge BEH column (Waters Corp, Milford, MA. 150mm x 2.1mm) was used for analysis. The following LC conditions were used for each run: Mobile phase A consisted of 0.1% formic acid in water, mobile phase B contained 0.1% formic acid in acetonitrile. A flow rate of 0.3 mL/minute and a column temperature of 30°C was used. The following gradient parameters were used during each run: 0-2 minutes at 25%B, 2-17 minutes going from 25% to 100%B, 17-22 minutes at 100%B, and an additional 10 minutes of re-equilibration at 25%B. 8-pt calibration curves were prepared in analyte stripped sera (Sigma, cat# SAE0012) on each day of batch analysis using PUFA/MUFA standards purchased from Cayman Chem (Cat#17941 and #17942). Concentrations ranged from 0.25ug/ml in the lowest calibrator to 50ug/ml for the highest calibrator. Stable isotope PUFA’s spiked into the extraction solution, as listed above, were used for peak normalization prior to calibration curve analysis. MS settings include: CUR:40, Ion spray voltage: 5500 in positive, −4500 in negative. CAD gas: 9, ion source temperature: 550°C, GS1: 40, GS2: 70. All other individual parameters (i.e. CE, DP, EP/CXP) are shown next to each compound in **Supplement S5**. Concentrations were back calculated from 8-pt calibration curves using Multiquant software (SCIEX Corp). Concentrations in liver tissue were normalized to tissue mass.

### LC-MS analysis of free fatty acids in Dgat2 ASO plasma samples

A mixture of 12:0-d23,14:0-d27,15:0-d29,16:0-d31,18:0-d35,18:1-d17,18:2-d11,20:4-d11,20:5-d5, 22:6-d5, and 24:0-d47 internal standards (Cayman) was added to 50uL of the homogenate or 20uL plasma. Fatty acids were extracted by a bi-phasic solution of acidified methanol and isooctane, derivatized using PFBB, and analyzed by GC-MS on an Agilent 6890N gas chromatograph equipped with an Agilent 7683 autosampler. Fatty Acids were separated using a 15m ZB-1 column (Phenomenex) and monitored using SIM identification. Analysis was performed using MassHunter software. The overall operation was modified from a previously established methodology(70).

### LC-MS analysis of phospholipids

A semi-targeted LC-MS/MS method was used for the profiling of phosphatidylcholines (PC), phosphatidylethanolamines (PE), lyso-PC’s and lyso-PE’s. LC-MS/MS conditions were adapted from previous literature, using the C18 X-bridge BEH column (71). Mobile phase A consisted of 50% ACN/0.1%FA and 2mM ammonium formate. Mobile phase B consisted of 90% isopropanol/10%ACN/0.1% FA and 2mM ammonium formate. The following gradient parameters were used during each run: 0-2 minutes at 25%B, 2-15 minutes going from 25% to 100%B, 15-22 minutes at 100%B, and an additional 15 minutes of re-equilibration at 25%B. The flow rate was 0.2 ml/min with column oven set to 38°C. All ions were monitored using positive ionization mode using similar MS settings as described above. The characteristic 184 fragment ion was used as the q3 mass of each PC precursor ion (q1) specified in the method, while the neutral loss of 141 was used to calculate the product ion of each PE as reported previously(72). Resolution of different saturation states and chain lengths were extensively tested using lyso-phosphatidylcholine, phosphatidylcholine, lyso-phosphatidylethanolamine, and phosphatidylethanolamine mixes (Cayman Chem. Cat#’s: 24331, 34800, 25844, and 24332 respectively). Peak areas of each phospholipid form were normalized to appropriate stable isotope internal standards using Multiquant software (SCIEX Corp).

### LC-MS analysis of glycerophospholipids

Glycerophospholipids were measured according to the method described previously with slight modifications(73,74). The Vanquish UHPLC (Thermo Fisher Scientific) was interfaced with a Q Exactive mass spectrometer (Thermo Fisher Scientific). Glycerolipids were analyzed using a data dependent acquisition (DDA) Top N scan of 8 with an NCE of 25 in positive mode. All ions in the mass range of 200-1200 m/z were monitored. MS1 resolution was set at 70,000 (FWHM at m/z 200) with an automatic gain control (AGC) target of 1e6 and Maximum IT of 200 ms. MS2 resolution was set to 17,500 with an AGC target of 5e4, fixed first mass of 80 m/z, and Maximum IT of 50 ms. The isolation width was set at 1.2; Dynamic Exclusion was set at 3 sec. Lipid Identification and Quantification was done with Lipidomic Data Analyzer Software(73,74).

### Digital Gene Expression (DGE) data generation from RNA-Seq Pseudo-Alignment and gene filtering

Salmon 0.7 was used per the standard protocol of single-end pseudo-alignment with default parameters. GRCm39 from GenBank was used as the reference genome. TPM was chosen as the quantification method for genes (75,76), and the quality of pseudo-alignment was checked both qualitatively and quantitatively. Briefly, to identify the best fit models for correcting heteroscedasticity, empirical CDF was visualized by five equally sized bins of different read lengths using different fit models (linear, Poisson, negative binomial with/without local adjustment). The goodness-of-fit for variance regression between observed and estimated variances was visually compared across these models, and the best model was selected based on their precision in estimating variances. A negative binomial model was assumed as the linkage type between variances and means when the p-values were computed for all unpaired hypothesis testing in gene differential analyses.

The TPM cutoff threshold for genes was determined holistically by visualizing the maintenance of minimal Pearson coefficient between within-group samples and the number of genes left after applying different cutoff thresholds. The minimal Pearson coefficient was selected from those computed between the two vectors of gene TPMs under a current threshold for each pair of samples within each condition group (e.g. Lean at week 1). Eventually, TPM threshold of >10, profiled-wide variance ranking among the top 70%, and unadjusted p values <0.5 were used as criteria to filter out genes. Among all pairwise differential gene expression analyses, approximately 4000-6000 genes were selected for further analysis out of 39538 after applying the criteria. Benjamini-Hochberg (BH) Procedure was used for FDR correction.

### Aggregated model fluxes for each sample used for machine learning

For each fitted model, an aggregate network was generated from all the fitted lipid models associated with the sample. The aggregated pseudo-fluxes Ϝ of all the reactions of each fitted model *z* of a sample *j*_1_ *was* computed as:

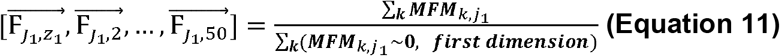

### Feature filtering with Minimum Redundancy Maximum Relevance (MRMR) Algorithm and multiclass classification by binary decision trees

Prior to machine learning, samples from the different treatment groups were partitioned with stratification, where 75% of the samples were used as the training set and 25% as the prediction set. The associated models of each sample were partitioned accordingly and used for the actual training and prediction. Aggregated models fluxes 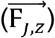 were treated as features and used for the prediction of treatment labels by classification trees (lean samples at week 1 & week 4, control samples at week 4, and *Dgat2* samples at week 4).

Prior to model training, MRMR was deployed to filter the features due to its excellent performance to identify mutually and maximally dissimilar features by exhaustively comparing pairwise features based on mutual information, which is intrinsically stable (77). These characteristics allowed us to identify the most representative aggregated model fluxes for each treatment group while addressing their co-dependency prior to applying classification. Features with the highest predictor scores (s c o re > 3\**St. dev*(all non-zero scores) or top 5% by values, whichever was bigger) and with all 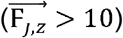 were kept for classification training and prediction.

Using the training set, a binary classification tree was trained with curvature test(78,79) as the predictor selection method, leave-one-out cross-validation, and an acquisition function described by Gelbart *et al* (80) for automatic hyperparameter optimization. Multiple trees were fitted with the training set, and the first tree with total misclassification rate <10% was chosen to predict the prediction set labels (maximum 20 attempts). The predicted label of a biological sample was then assigned by the most frequently predicted labels among the 50 associated models. Parameter specifications for training were inputted in conjunction with the *fitctree* function in MATLAB.

To test the robustness and accuracy of the machine learning method, the process was repeated 100 times by randomly partitioning samples into training and prediction sets, and the overall performance of the method was evaluated by the ratio of correctly predicted labels of the prediction set (sensitivity).

### Statistical Tests

If not clearly specified, comparison of model pseudo-fluxes or pseudo-concentrations (including fold changes) was conducted by Wilcoxon’s rank sum test with a significance (p-value) threshold of 0.01. For comparison of gene expressions, unpaired, heteroscedastic Student’s t-test was conducted with a significance (p-value) threshold of 0.05.

## Notes

### Competing Interest Statement

The authors have declared no competing interest.

### Summary of Updates

The coauthor name "Andrey Low" has been changed to "Audrey Low"

