## Supplemental for "LipidSIM: inferring mechanistic lipid biosynthesis perturbations from lipidomics with a flexible, low-parameter, systematic Markov Modeling framework"

### Supplement

#### *S1 Reaction rules for de novo lipid synthesis network construction*

*k: reaction index; n: number of carbons; x: number of unsaturated bonds

**: While reaction rules for sterol and ether lipids were also curated, they were not included in the models used for this study due to lack of relevant lipidomic data.

| Type | Primary Gene/Enzyme/Rxn Symbol/Abbreviations | Reactant(s) | Product | Constraints |
|---|---|---|---|---|
| Saturated FA | FASN_1 | Malonyl-CoA;Acetyl-CoA | 4:0-CoA | |
| Saturated FA | FASN_2 | Malonyl-CoA;Propionyl-CoA | 5:0-CoA | |
| Multi-unsaturated | FADS2_1 | 9,12-18:2-CoA | 6,9,12-18:3-CoA | |
| Multi-unsaturated | FADS1_2 | 8,11,14-20:3-CoA | 5,8,11,14-20:4-CoA | |
| Multi-unsaturated | FADS2_3 | 7,10,13,16-22:4-CoA | 4,7,10,13,16-22:5-CoA | |
| Multi-unsaturated | FADS2_4 | 9,12,15,18-24:4-CoA | 6,9,12,15,18-24:5-CoA | |
| Multi-unsaturated | FADS2_5 | 9,12,15-18:3-CoA | 6,9,12,15-18:4-CoA | |
| Multi-unsaturated | FADS1_6 | 8,11,14,17-20:4-CoA | 5,8,11,14,17-20:5-CoA | |
| Multi-unsaturated | FADS2_7 | 7,10,13,16,19-22:5-CoA | 4,7,10,13,16,19-22:6-CoA | |
| Multi-unsaturated | BOxi_8 | 6,9,12,15,18,21-24:6-CoA | 4,7,10,13,16,19-22:6-CoA | |
| Multi-unsaturated | BOxi_9 | 6,9,12,15,18-24:5-CoA | 4,7,10,13,16-22:5-CoA | |
| Multi-unsaturated | FADS2_9 | 9,12,15,18,21-24:5-CoA | 6,9,12,15,18,21-24:6-CoA | |
| VLCFA | ELOVL_1 | 6,9,12-18:3-CoA;Acetyl-CoA | 8,11,14-20:3-CoA | |
| VLCFA | ELOVL_2 | 5,8,11,14-20:4-CoA;Acetyl-CoA | 7,10,13,16-22:4-CoA | |
| VLCFA | ELOVL_3 | 6,9,12,15-18:4-CoA;Acetyl-CoA | 8,11,14,17-20:4-CoA | |
| VLCFA | ELOVL_4 | 5,8,11,14,17-20:5-CoA;Acetyl-CoA | 7,10,13,16,19-22:5-CoA | |
| VLCFA | ELOVL_5 | 7,10,13,16,19-22:5-CoA;Acetyl-CoA | 9,12,15,18,21-24:5-CoA | |
| VLCFA | ELOVL_8 | 7,10,13,16-22:4-CoA;Acetyl-CoA | 9,12,15,18-24:4-CoA | |
| Sterol | HSD3B | Cholesterol | Cholestenone | |
| Sterol | HMGCS;HMGCR; MVK;PMVK;MVD | Acetyl-CoA;Acetoacetyl-CoA | 3,3-dimethylallylpyrophosphoric acid | |
| Sterol | IDI1 | 3,3-dimethylallylpyrophosphoric acid | 5-isopentenylpyrophosphoric acid | |
| Sterol | FDPS;GGPS;FDFT;SQLE;LSS | 3,3-dimethylallylpyrophosphoric acid; 5-isopentenylpyrophosphoric acid | Lanosterol | |
| Sterol | Cyp51_1;LBR_1;NSDHL_1 | Lanosterol | 24-dehydrolathosterol | |
| Sterol | SC5D_1 | 24-dehydrolathosterol | 7-dehydrodesmosterol | |
| Sterol | DHCR7_1 | 7-dehydrodesmosterol | Desmosterol | |
| Sterol | DHCR24_1 | Lanosterol | Cholesta-8(9)-en-3B-ol | |
| Sterol | Cyp51_2;LBR_2;NSDHL_2 | Cholesta-8(9)-en-3B-ol | Lathosterol | |
| Sterol | SC5D_2 | Lathosterol | 7-dehydrocholesterol | |
| Sterol | DHCR7_2 | 7-dehydrocholesterol | Cholesterol | |
| Sterol | DHCR24_2 | Desmosterol | Cholesterol | |
| Sterol | Cyp7A1 | Cholesterol | 7alpha-hydroxy-cholesterol | |
| Sterol | Cyp3A4 | Cholesterol | 4beta-hydroxy-cholesterol | |
| Sterol | Cyp46A1 | Cholesterol | 24S-hydroxy-cholesterol | |
| Sterol | CH25H | Cholesterol | 25-hydroxy-cholesterol | |
| Sterol | CYP27A1_1 | Cholesterol | 27-hydroxy-cholesterol | |

| | | | | |
|---|---|---|---|---|
| Sterol | ROS_1 | Cholesterol | 7-oxo-cholesterol | |
| Sterol | LCAT | 7-oxo-cholesterol | 7-oxo-cholesterol ester (PC variety) | |
| Sterol | SOAT1/2 | 7-oxo-cholesterol | 7-oxo-cholesterol ester (FA variety) | |
| Sterol | Cyp27A1_2 | 7-oxo-cholesterol | 27-hydroxy-7-oxo-cholesterol | |
| Sterol | HSD11B1 | 7-oxo-cholesterol | 7beta-hydroxy-cholesterol | |
| Saturated FA | FASN_k | n:0-CoA;Acetyl-CoA | (n+2):0-CoA | 4<= n <=18 |
| Mono-unsaturated FA | MUFADS2_k | n:0-CoA | (n-9)-n:1-CoA | 14<= n <=19 |
| Eicosanoids | COX_1 | 8,11,14-20:3-CoA | PG1 | |
| Eicosanoids | COX_2 | 5,8,11,14-20:4-CoA | PG2 | |
| Eicosanoids | COX_3 | 5,8,11,14,17-20:5-CoA | PG3 | |
| PA | GPAT_k | n:x-CoA | LPA(n1:x1) | 14<=n<=24, x<=5 |
| PA | AGPAT_k | LPA(n1:x1);…-n2:x2-CoA | PA(n1+n2:x1+x2) | 14<=n<=24, x<=5, abs(n1-n2)<=4 |
| Diacylglycerol | PAP_k | PA(n1+n2:x1+x2) | 1_2-DG(n1+n2:x1+x2) | 14<=n<=24, x<=5, abs(n1-n2)<=4 |
| Diacylglycerol | DAG_isomerization_k | 1_2-DG(n1+n2:x1+x2) | 1_3-DG(n1+n2:x1+x2) | |
| Triacylglycerol | DGAT_k | 1_2-DG(n1+n2:x1+x2) | TG(n1+n2+n3:x1+x2+x3) | |
| Sterol | ACAT | Cholesterol;…-n:x-CoA | CE(n1:x1) | 14<=n<=22,x<=4 |
| PC | CPT;PLC_k | 1_2-DAG(n1+n2:x1+x2) | PC(n1+n2:x1+x2) | 14<=n<=24 |
| PC | PEMT_k | PE(n1+n2:x1+x2) | PC(n1+n2:x1+x2) | 14<=n<=24 |
| PC | PLA2G6_1;LPCAT_1_k | PC(n1+n2:x1+x2) | sn1-LPC(n1:x1);sn2-LPC(n2:x2); | 14<=n<=24 |
| PA | PLD2_k | PC(n1+n2:x1+x2) | PA(n1+n2:x1+x2) | 14<=n<=24 |
| PE | EPT_k | 1_2-DG(n1+n2:x1+x2) | PE(n1+n2:x1+x2) | 14<=n<=24 |
| PE | PLA2G6_2;LPCAT_2_k | PE(n1+n2:x1+x2) | sn1-LPE(n1:x1);sn2-LPE(n2:x2); | 14<=n<=24 |
| PG | CDS1;PGS;PTPMT_k | PA(n1+n2:x1+x2);G3P | PG(n1+n2:x1+x2) | 14<=n<=24 |
| PI | CDIPT_k | PA(n1+n2:x1+x2) | PI(n1+n2:x1+x2) | 14<=n<=24 |
| PS | CDS1;CHO1_k | PA(n1+n2:x1+x2) | PS(n1+n2:x1+x2) | 14<=n<=24 |
| PS | PISD/PTDSS2_k | PS(n1+n2:x1+x2) | PE(n1+n2:x1+x2) | 14<=n<=24 |
| PS | PTDSS1_k | PC(n1+n2:x1+x2) | PS(n1+n2:x1+x2) | 14<=n<=24 |
| Ether Lipid | GNPAT;AGPS;R03455(etc)_k | G3P;...n:x-CoA | 1-O-alkyl(n:x) | n=16,18,20,22 |
| Ether Lipid | EPT_like_k | ...n2:x2-CoA;1-O-alkyl(n1:x1) | PE(n1:x1) | x<=7, abs(n1-n2)<=4 |
| Ether Lipid | CPT_like_k | ...n2:x2-CoA;1-O-alkyl(n1:x1) | PC(n1x1) | x<=7, abs(n1-n2)<=4 |
| Ether Lipid | PEDS1_1_k | PE(n1+n2:x1+x2e) | PE(n1:x1) | |
| Ether Lipid | PEDS1_2_k | PC(n1+n2:x1+x2e) | PC(n1:x1) | |
| Ether Lipid | PLA2G6_3;LPCAT_3_ | PE(n1+n2:x1+x2e) | LPE(n1:x1e) | |
| Ether Lipid | PLA2G6_4;LPCAT_4_ | PC(n1+n2:x1+x2e) | LPC(n1:x1e) | |
| Ether Lipid | PLA2G6_5;LPCAT_5_ | PE(n1+n2:x1+x2p) | LPE(n1:x1p) | |
| Ether Lipid | PLA2G6_6;LPCAT_6_ | PC(n1+n2:x1+x2p) | LPC(n1:x1p) | |

#### S2 Volcano plot for differential fatty acid species between KEAP1 ASO treated samples and control ASO treated samples

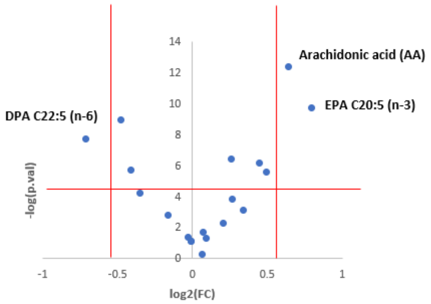


#### S3 Aggregated pseudo-concentrations of lipid species by the number of unsaturated carbon bonds and the number of acyl carbons for control or KEAP1 ASO treatment groups from KEAP1 dataset

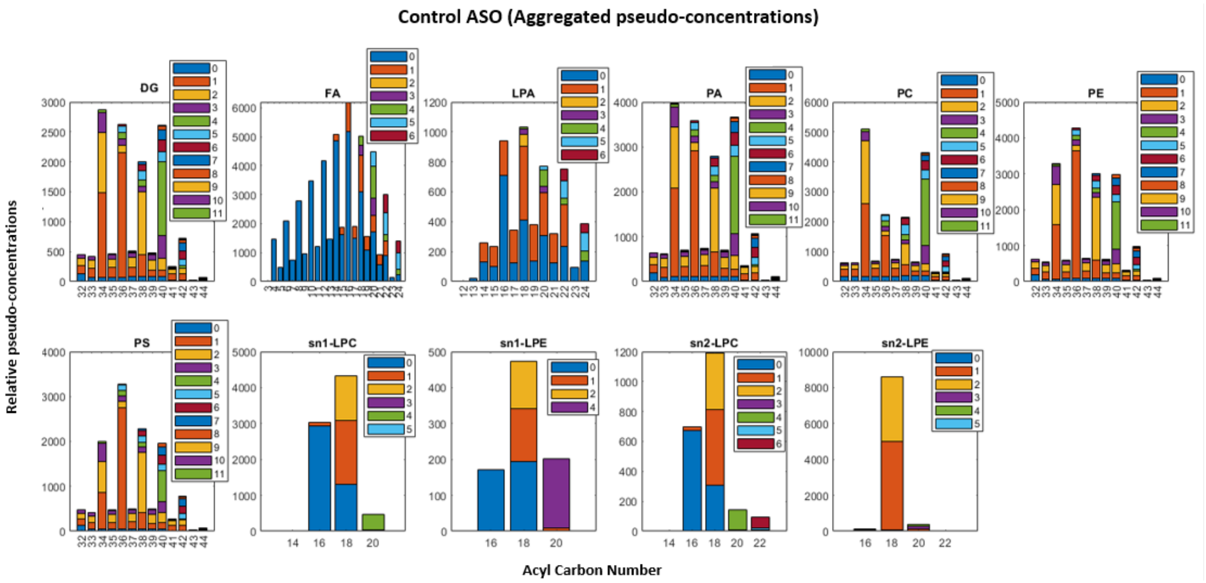

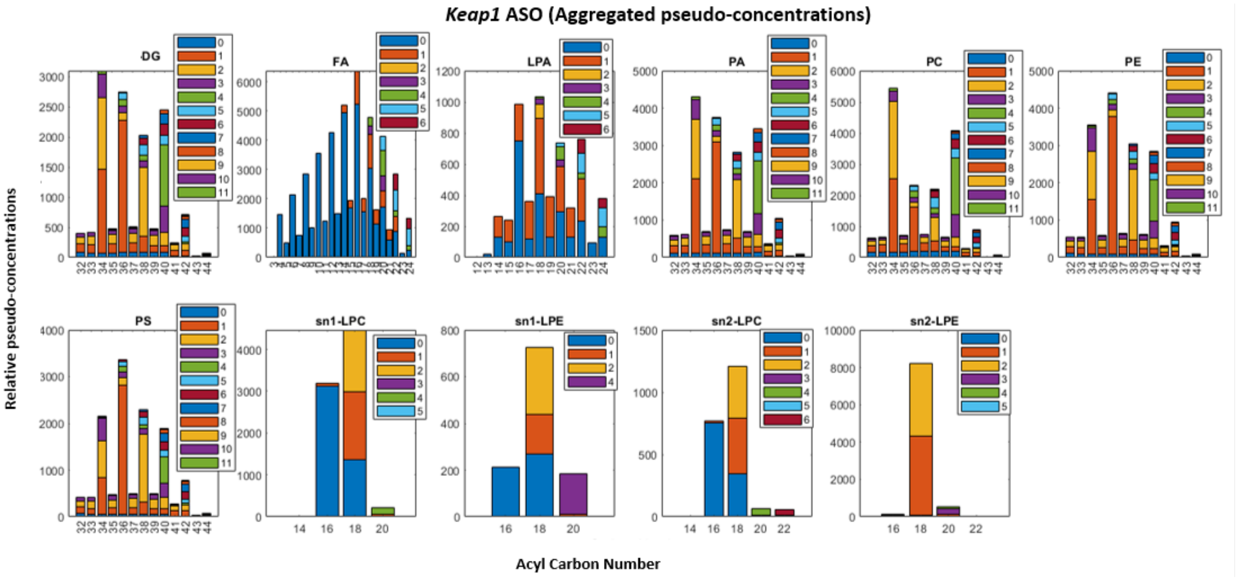


#### *S4. Top splitting nodes based on their frequencies from 100 fitted decision trees*

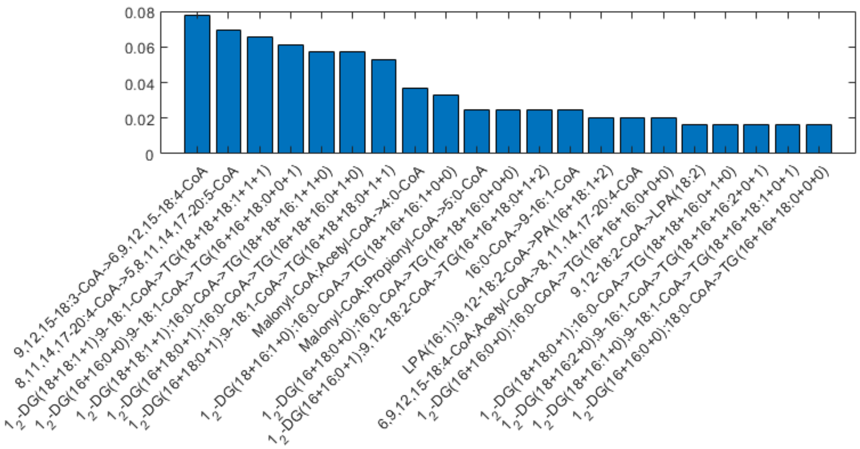


#### *S5. Compound list and parameter table for positive and negative ionization polarity switching method*

MRM transition (Q1 and Q3 m/z), declustering potential, collision energy, Entrance potential (EP), and CXP (collision cell exit potential) are detailed. In addition, intra-batch precision results (from 2 levels of QC's, 150ng/ml and 800ng/ml final plasma concentrations, respectively) and accuracies are shown for compounds where calibration curves were run. The first transition for each compound is used for quantitation while the second was used as confirmation. Accuracy and precision are not shown for several compounds because of the natural presence of these compounds in the matrix used (analyte stripped sera) causing interference with these calculations for the low QC.

| Compound | Q1 m/z | Q3 m/z | Declustering potential (DP) | Collision energy (CE) | EP | CXP | Ionization mode | %CV (Avg. Low, high) | %Accuracy (Avg. low, high) |
|---|---|---|---|---|---|---|---|---|---|
| Myrsitoleic C14:1 | 227.2 | 191.1 | 60 | 13 | 7 | 7 | POS | | |
| Myrsitoleic C14:1 2 | 227.2 | 227.2 | 60 | 10 | 7 | 7 | POS | | |
| Palmitoleic C16:1 | 255.2 | 219.2 | 60 | 13 | 7 | 7 | POS | 9, 9 | 115, 103 |
| Palmitoleic C16:1 2 | 255.2 | 97.1 | 60 | 22 | 7 | 7 | POS | | |
| Oleic acid C18:1 | 283.3 | 247.2 | 60 | 12 | 7 | 7 | POS | | |
| Oleic acid C18:1 2 | 283.3 | 97.1 | 60 | 25 | 7 | 7 | POS | | |
| Linoleic C18:2 | 281.3 | 245 | 70 | 15 | 5 | 7 | POS | 18, 8 | 81, 95 |
| Linoleic C18:2 2 | 281.3 | 55 | 70 | 57 | 5 | 7 | POS | | |
| d4 Linoleic | 285.3 | 249 | 70 | 15 | 5 | 7 | POS | | |
| Linolenic C18:3 2 | 279.2 | 67 | 70 | 50 | 5 | 7 | POS | | |
| Linolenic C18:3 3 | 279.2 | 81 | 70 | 32 | 5 | 7 | POS | | |
| Stearidonic C18:4 1 | 277.2 | 79 | 70 | 55 | 10 | 7 | POS | 7, 3 | 99, 91 |
| Stearidonic C18:4 2 | 277.2 | 121 | 70 | 27 | 10 | 7 | POS | | |
| Dihomolinoleic C20:3 1 | 307.3 | 67 | 60 | 55 | 10 | 7 | POS | | |
| Dihomolinoleic C20:3 2 | 307.3 | 81 | 60 | 43 | 10 | 7 | POS | | |
| Eicosapentaenoic acid C20:5 | 303.2 | 91 | 70 | 57 | 5 | 7 | POS | 6, 7 | 99, 95 |
| Eicosapentaenoic acid C20:5 2 | 303.2 | 81 | 70 | 42 | 5 | 7 | POS | | |
| Arachidonic acid C20:4 1 | 305.3 | 93 | 70 | 42 | 10 | 7 | POS | 6, 4 | 87, 97 |
| Arachidonic acid C20:4 2 | 305.3 | 221.2 | 60 | 19 | 10 | 7 | POS | | |
| Adrenic acid C22:4 1 | 331.3 | 287.2 | -90 | -23 | -13 | -9 | NEG | 7, 2 | 108, 95 |
| Adrenic acid C22:4 2 | 331.3 | 233.3 | -90 | -24 | -13 | -9 | NEG | | |
| Docosapentaenoic acid C22:5 1 | 329.2 | 285.2 | -90 | -20 | -13 | -9 | NEG | 9, 9 | 94, 102 |
| Docosapentaenoic acid C22:5 2 | 329.2 | 231.1 | -90 | -22 | -13 | -9 | NEG | | |
| Docosahexanoic C22:6 1 | 327.2 | 283.2 | -90 | -18 | -13 | -9 | NEG | 5, 12 | 108, 106 |
| Docosahexanoic C22:6 2 | 327.2 | 229.2 | -90 | -18 | -13 | -9 | NEG | | |
| 12-OH Lauric acid | 215.2 | 169.1 | -80 | -25 | -7 | -5 | NEG | 9, 16 | 101, 90 |
| 12-OH Lauric 2 | 215.2 | 215.2 | -80 | -10 | -7 | -5 | NEG | | |
| 13-HOTrE 1 | 293.2 | 195.2 | -85 | -22 | -7 | -5 | NEG | 9, 14 | 92, 93 |
| 13-HOTrE 2 | 293.2 | 223 | -85 | -23 | -7 | -5 | NEG | | |
| 11-HEDE | 323.3 | 199 | -90 | -26 | -13 | -8 | NEG | 8, 13 | 104, 102 |
| 11-HEDE 2 | 323.3 | 305 | -90 | -27 | -13 | -8 | NEG | | |
| 15-HETE | 319.2 | 219.1 | -75 | -18 | -7 | -9 | NEG | 15, 15 | 93, 102 |
| 12-HETE | 319.2 | 179.1 | -75 | -20 | -7 | -9 | NEG | 4, 9 | 105, 119 |
| 11-HETE | 319.2 | 167.1 | -75 | -20 | -7 | -9 | NEG | | |
| 5-HETE | 319.2 | 115 | -75 | -17 | -7 | -9 | NEG | 17, 10 | 114, 106 |
| PGE2_1 | 351.2 | 271.2 | -65 | -22 | -10 | -10 | NEG | 9, 6 | 93, 104 |
| PGE2_2 | 351.2 | 315.2 | -65 | -15 | -10 | -10 | NEG | | |
| D4-PGE2 | 355.3 | 275.2 | -65 | -22 | -10 | -10 | NEG | | |
| d8-arachidonic acid | 311.3 | 267.2 | -90 | -20 | -13 | -9 | NEG | | |
